# The SARS-CoV-2 and other human coronavirus spike proteins are fine-tuned towards temperature and proteases of the human airways

**DOI:** 10.1101/2020.11.09.374603

**Authors:** Manon Laporte, Annelies Stevaert, Valerie Raeymaekers, Ria Van Berwaer, Katleen Martens, Stefan Pöhlmann, Lieve Naesens

## Abstract

The high transmissibility of SARS-CoV-2 is related to abundant replication in the upper airways, which is not observed for the other highly pathogenic coronaviruses SARS-CoV-1 and MERS-CoV. We here reveal features of the coronavirus spike (S) protein, which optimize the virus towards different parts of the respiratory tract. First, the SARS-CoV-2 spike (SARS-2-S) reached higher levels in pseudoparticles when produced at 33°C instead of 37°C. Even stronger preference for the upper airway temperature of 33°C was evident for the S protein of HCoV-229E, a common cold coronavirus. In contrast, the S proteins of SARS-CoV-1 and MERS-CoV favored 37°C, in accordance with their preference for the lower airways. Next, SARS-2-S proved efficiently activated by TMPRSS13, besides the previously identified host cell protease TMPRSS2, which may broaden the cell tropism of SARS-CoV-2. TMPRSS13 was found to be an effective spike activator for the virulent coronaviruses but not the common cold HCoV-229E virus. Activation by these proteases requires pre-cleavage of the SARS-2-S S1/S2 cleavage loop, and both its furin motif and extended loop length proved critical to achieve virus entry into airway epithelial cells. Finally, we show that the D614G mutation in SARS-2-S increases S protein stability and expression at 37°C, and promotes virus entry via cathepsin B/L activation. These spike properties might promote virus spread, potentially explaining why the G614 variant is currently predominating worldwide. Collectively, our findings indicate how the coronavirus spike protein is fine-tuned towards the temperature and protease conditions of the airways, to enhance virus transmission and pathology.

**SIGNIFICANCE STATEMENT:** The rapid spread of SARS-CoV-2, the cause of COVID-19, is related to abundant replication in the upper airways, which is not observed for other highly pathogenic human coronaviruses. We here reveal features of the coronavirus spike (S) protein, which optimize the virus towards different parts of the respiratory tract. Coronavirus spikes exhibit distinct temperature preference to precisely match the upper (~33°C) or lower (37°C) airways. We identified airway proteases that activate the spike for virus entry into cells, including one protease that may mediate coronavirus virulence. Also, a link was seen between spike stability and entry via endosomal proteases. This mechanism of spike fine-tuning could explain why the SARS-CoV-2 spike-D614G mutant is more transmissible and therefore globally predominant.

## INTRODUCTION

The devastating COVID-19 pandemic is caused by the novel bat-derived SARS-CoV-2 virus. Despite its recent introduction in humans, this coronavirus (CoV) is already very well adapted for efficient respiratory droplet transmission and high-titer replication in human airways (1, 2). Its disease spectrum varies from mild respiratory symptoms to severe pneumonia (3), depending mostly on the patient’s age and comorbidities. The SARS-CoV-2 pandemic was preceded by local outbreaks of SARS-CoV-1 and MERS-CoV, two other highly virulent viruses with a bat origin (4, 5). Compared to SARS-CoV-2, SARS-CoV-1 and MERS-CoV have far lower tropism for the upper respiratory tract (1, 6). In contrast, a mild common cold-like disease is typical for the endemic human CoVs 229E, NL63, OC43 and HKU1 (7). Their zoonotic spillover probably occurred long time ago (8–10), implying extensive adaptation to the upper respiratory tract in which these common cold viruses are flourishing.

CoV replication in the upper or lower airways implicates that the virus is compatible with the temperature in these compartments, which evolves from ~30-32°C in the nose to 37°C in the deeper airways (11, 12). We recently showed that the hemagglutinin of influenza B virus has an intrinsic preference for 33°C to be robustly expressed, consistent with 33°C being the best temperature to propagate this virus. Other temperature profiles were recognized for the hemagglutinin proteins of human and avian influenza A viruses (13). This subtle adaptation of viral glycoproteins to the temperature in the host organs might also apply to other respiratory viruses with a zoonotic origin, in particular CoVs. Since SARS-CoV-2 exhibits abundant replication in the nose (14), it is conceivable that its spike (S) glycoprotein (SARS-2-S) is fine-tuned towards this cooler compartment.

The trimeric spike protein mediates CoV entry into the host cell. Its S1 domain is responsible for receptor binding, while the S2 domain mediates fusion between the viral envelope and a cellular membrane (15). To become membrane fusion-competent, the full-length spike protein (S0) needs to be cleaved by host cell proteases at its S1/S2 and S2’ sites (16, 17). Cleavage at the S2’ site might be sufficient to trigger membrane fusion and is subsequently referred to as activation, since it releases the internal fusion peptide (18). The host protease TMPRSS2 is a prominent player in S2’ activation, however also other proteases may be involved, thus broadening the cell or tissue tropism of SARS-CoV-2. TMPRSS2 knockout was shown to reduce lung pathology in mouse models for SARS and MERS (19). Still, the finding that virus replication was slower but not abolished, suggests that activation can be performed by additional proteases. The Type II Transmembrane Serine Protease (TTSP) family, to which TMPRSS2 belongs, contains in total 18 proteases, many of which are expressed in human airways (13). Only a limited number have been investigated in the CoV context [reviewed in: (16)]. Precise knowledge of the activating TTSPs would help to conceive host-directed therapies interfering with spike activation (20).

One noticeable feature of the SARS-CoV-2 spike is the presence of an extended S1/S2 cleavage loop bearing a multibasic furin recognition motif (RRAR) (21). This loop extension is the result of a four-amino acid-insertion not present in the S protein of the probable bat ancestor virus (22). Pre-cleavage of the multibasic motif is assumedly performed by furin-like proteases (23–26), but also other proteases have been proposed (27). S1/S2 pre-cleavage proved crucial for subsequent SARS-2-S activation by TMPRSS2 and for viral entry into airway epithelial Calu-3 cells (24). The furin dependence also applies to the S protein of MERS-CoV (MERS-S) (28, 29), but not the spike protein of SARS-CoV-1 (SARS-1-S) (24). On the other hand, all three aforementioned CoV S proteins do not need S1/S2 pre-cleavage to mediate entry into cells expressing the endo/lysosomal protease cathepsin B/L, which activates the S protein after virus uptake by endocytosis (18, 25). For SARS-2-S, the determinants governing cleavage of its extended S1/S2 loop are still far from clear. In addition, SARS-CoV-2 passaging in Vero cells commonly leads to substitutions or deletions in the S1/S2 cleavage loop (30–36). In animal models, these viruses exhibit reduced transmission (37) or virulence (38). Mutant viruses bearing substitutions or deletions in the S1/S2 loop are rarely detected in humans (33, 37, 39) (Suppl. Tables S1 and S2).

Within a few months circulation among humans, SARS-CoV-2 has acquired a mutation (D614G) in its spike protein, that is increasingly recognized as a possible cause of enhanced virus transmission (40–42). The D614G mutation was already detected during the early phase of the pandemic and, after four months, the S^G614^ variant became globally predominant (43). Compared to SARS-CoV-2-S^D614^ virus, the S^G614^ variant exhibits higher transmissibility in animal models (40–42). The D614G mutation was also shown to enhance SARS-CoV-2 replication (40–42) and increase infectivity of SARS-2-S-bearing pseudoparticles (40, 42, 44–47). Hence, both the D614G mutation and S1/S2 cleavage loop deletions appear to alter virus transmissibility, but the underlying mechanisms remain enigmatic.

The aim of this study was to assess how SARS-2-S is fine-tuned towards the temperature and proteases of the airways; and how these properties compare to those of SARS-1-S, MERS-S and the S protein of the common cold virus HCoV-229E. By performing pseudovirus production at 33°C and 37°C, we revealed that each spike protein exhibits an intrinsic and distinct temperature preference, correlating with viral preference for the upper or lower airways. We next addressed how SARS-2-S driven entry is controlled by host proteases that cleave its extended S1/S2 loop or S2’ site. Hence, we studied the entry behavior of different SARS-2-S loop deletion mutants, and we assessed which of the 18 human TTSPs act as CoV spike activators. Finally, we investigated the temperature and protease dependency of the SARS-2-S D614G mutant, to appreciate how these spike features might be linked to virus transmissibility.

## RESULTS

### The spikes of SARS-CoV-2 and HCoV-229E prefer 33°C for pseudoparticle production, while the SARS-CoV-1 spike prefers 37°C

The temperature gradient in the human respiratory tract is a plausible key factor in determining whether a virus preferentially replicates in the upper or lower airways. We observed that the common cold virus HCoV-229E replicates more efficiently at 33°C and 35°C, when compared to 37°C and, particularly, 39°C (Suppl. Fig. 1). Similarly, SARS-CoV-2 was reported to prefer 33°C to 37°C, while SARS-CoV-1 showed the opposite profile (48).

We hypothesized that CoV S proteins may exhibit temperature dependency, in relation to the temperature required for optimal virus replication. To study this, we produced MLV particles bearing the S proteins of the highly pathogenic SARS-CoV-1 (SARS-1-S), SARS-CoV-2 (SARS-2-S) and MERS-CoV (MERS-S) or the common cold virus HCoV-229E (229E-S) (Fig. 1A). Immunoblot analysis of pseudoparticles produced at 33°C or 37°C revealed that temperature impacted particle incorporation of these S proteins, but not S protein cleavage by host cell proteases. Thus, particle incorporation of SARS-2-S^D614^ was 1.9-fold (P=0.027) higher at 33°C than at 37°C (Fig. 1B). This effect was even more pronounced for 229E-S (= 10-fold higher level at 33°C than at 37°C; P=0.0027), which was barely incorporated at 37°C. The picture was entirely opposite for SARS-1-S, where particle incorporation was 2.7-fold higher at 37°C than at 33°C (P=0.0047) and a similar trend was observed for MERS-S.

**Figure 1.**
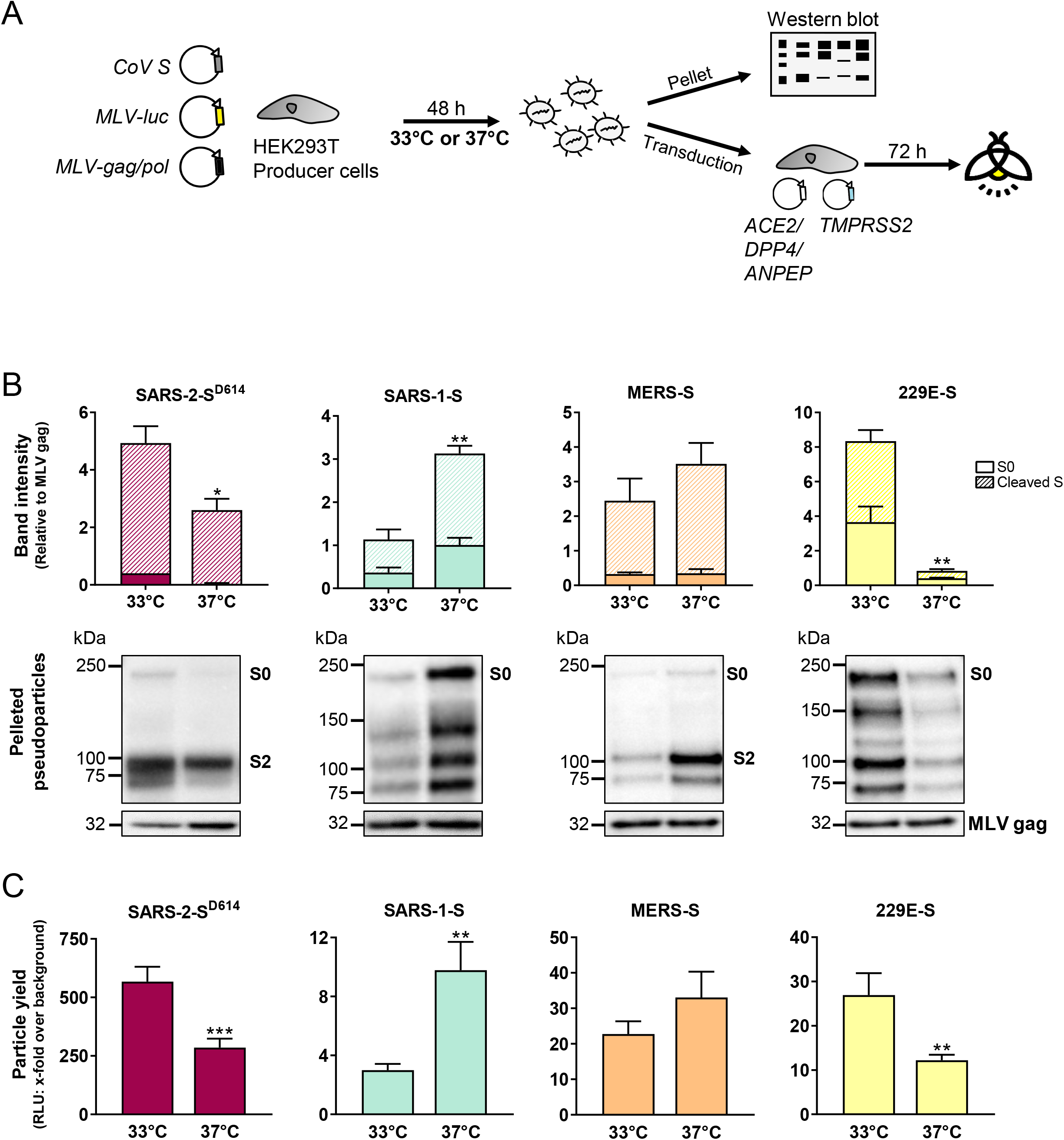
Spike incorporation into pseudovirions is temperature-dependent. (A) Experiment set-up. S-bearing pseudoviruses were produced in HEK293T cells at either 33°C or 37°C, and the released particles were pelleted to determine S content by western blot. In parallel, they were used to transduce HEK293T target cells expressing the appropriate receptor and TMPRSS2. (B) The graphs show S content relative to that of MLV-gag (mean ± SEM; of three independently produced stocks). Representative blots show uncleaved and cleaved S protein bands. (C) Particle infectivity was measured by luminescence read-out at day 3 post transduction. A two-tailed unpaired t-test was used to compare the 33°C and 37°C results, regarding total S content or particle infectivity. *, P ≤ 0.05; **, P ≤ 0.01; ***, P ≤ 0.001.

Importantly, for each virus, the temperature preference was equally apparent when we determined particle infectivity by transducing HEK293T cells transfected with TMPRSS2 and the appropriate receptor [i.e. human ACE2 for SARS-CoV-1 and SARS-CoV-2; human dipeptidyl peptidase-4 (DPP4) for MERS-CoV; and human aminopeptidase N (APN) for HCoV-229E] (Fig. 1C). Pseudoparticles produced at 33°C harboured more SARS-2-S and 229E-S and showed higher infectivity as compared to their counterparts produced at 37°C, and the reverse effect was observed for particles bearing SARS-1-S or MERS-S. For each pseudovirus, the signal was the same whether virus entry (i.e. target cell transduction) was performed at 33°C or 37°C (data not shown).

Collectively, these results indicate that CoV S protein appearance in virus particles is temperature-dependent, in relation to the temperature required for optimal virus replication. The SARS-2-S^D614^ protein prefers 33°C to 37°C, a property that is still more pronounced for 229E-S. On the contrary, 37°C is the preferred temperature for MERS-S and especially SARS-1-S.

### Generation of S-pseudoviruses with wild-type and mutant spikes and mRNA expression analysis of target cells

To investigate the determinants and impact of S1/S2 pre-cleavage in SARS-2-S (Fig. 2A), we generated pseudoviruses with wild-type (WT) SARS-2-S^D614^ and three deletion mutants missing parts of the extended S1/S2 cleavage loop (Fig. 2B and 2C). The ΔPRRA mutant lacks the furin cleavage motif (RRAR) and its cleavage site is identical to that of the SARS-CoV-2-related bat CoV RaTG13 (22). The ΔQTQTN mutant lacks a sequence preceding the RRAR motif, while mutant ΔNSPRRAR lacks the RRAR motif plus three flanking amino acid residues. These or very similar deletions are commonly detected during passaging of SARS-CoV-2 in Vero cells (see Suppl. Table S3 and references therein), suggesting that they might confer a growth advantage in this cell line. For comparison, we introduced the multibasic cleavage site of SARS-2-S and its preceding residues into SARS-1-S, and we generated a MERS-S mutant in which the furin motif was destroyed (R748C). Besides these S1/S2 loop mutants, we produced the D614G mutant form of SARS-2-S, to investigate the possible mechanism behind the shift from the early S^D614^ to currently predominating S^G614^ variant virus (43). Residue 614 is located at an inter-protomer interface in the spike trimer (Fig. 2C) (44, 45).

**Figure 2.**
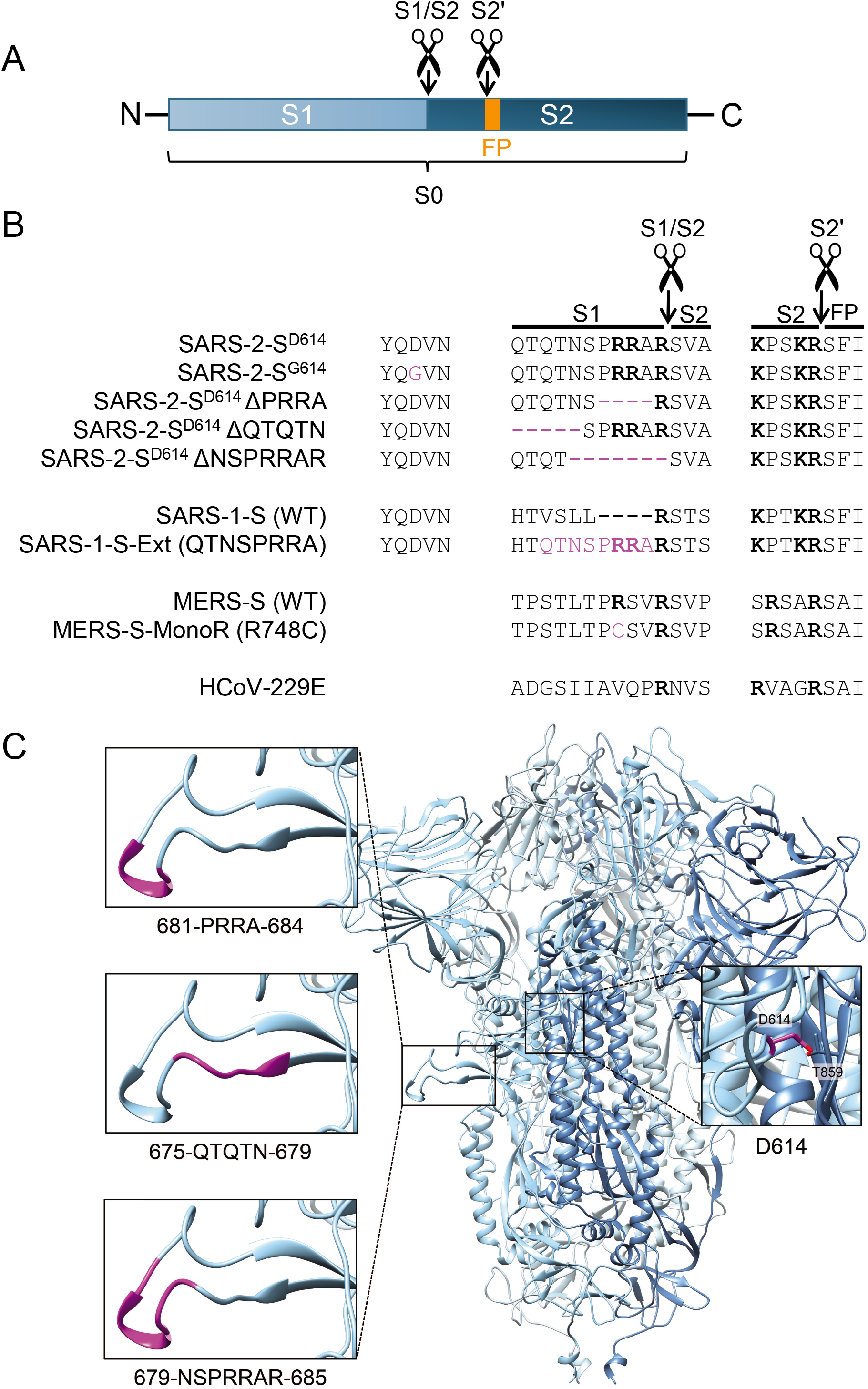
Generation of S-pseudoviruses with wild-type spikes, S1/S2 site mutant forms, and the D614G mutant of SARS-2-S. (A) The CoV S protein contains two main cleavage sites: the S1/S2 site separates the S1 and S2 subunits, whereas S2’ cleavage liberates the fusion peptide (FP). (B) Amino acid sequences around the S1/S2 and S2’ cleavage sites of the CoV spikes and mutant forms created in this study. Basic Arg and Lys residues are shown in bold. For SARS-2-S, the D614/G614 variation site is added. (C) Structure of the SARS-CoV-2 spike trimer, based on PDB 6ZGE (81), in which we modelled the cleavage loop using SWISS-MODEL (82). The amino acids shown in magenta were substituted or deleted, to create three S1/S2 loop mutants. The inset on the right shows residue D614, which forms a hydrogen bond with residue T859 in the S2 subunit of another protomer (44, 45).

Before conducting pseudovirus entry studies, we performed RT-qPCR analysis to estimate expression of relevant host cell factors in the three cell lines under study, namely human airway epithelial Calu-3 cells; African green monkey kidney Vero E6 cells; and human embryonic kidney HEK293T cells. Human nasal and lung tissue samples were included for comparison (Fig. 3A). Respiratory tissue contained the transcripts for three CoV receptors [i.e. ACE2, the entry receptor for SARS-CoV-1 and SARS-CoV-2; DPP4, the receptor for MERS-CoV; and APN, the HCoV-229E receptor], and the mRNA levels were comparable for the two anatomic sites. *ACE2* and *DPP4*, but not *ANPEP/APN*, were expressed in the three cell lines. Human nasal and lung tissue samples contained abundant mRNA levels for *CTSB* (cathepsin B) and intermediate levels for *CTSL* (cathepsin L). *CTSB* mRNA was also present in Vero E6 and HEK293T cells, and at lower levels in Calu-3 cells. The transcript levels of *CTSL* (cathepsin L) proved very high in Vero E6 cells, intermediate in HEK293T cells, and lower in Calu-3 cells, as described by others (28, 49). Finally, all samples contained quite comparable levels of *furin* mRNA, except for those of Calu-3, which were slightly lower. As reported by us before (13), Calu-3 cells show high expression of *TMPRSS2* and several other TTSPs, at levels comparable to those found in primary respiratory tissue. To conclude, Calu-3 cells are suitable to assess viral entry via TMPRSS2 and other TTSPs, while Vero E6 cells are useful to study cathepsin B/L-mediated entry. HEK293T cells are suited for ectopic expression of receptors or proteases.

**Figure 3.**
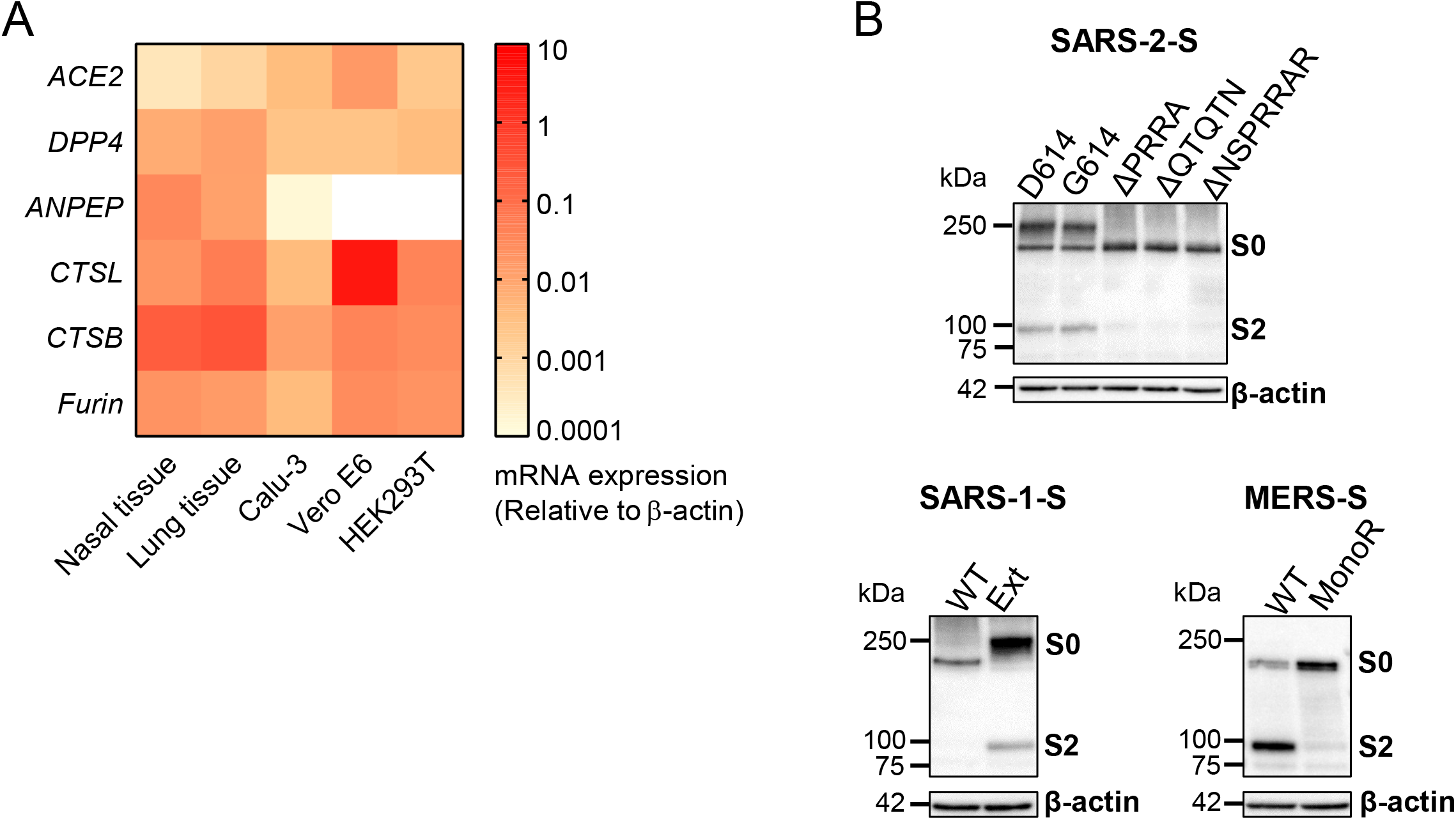
Expression of coronavirus receptors, activating proteases and spike proteins. (A) Expression of coronavirus receptors and activating proteases, determined by qRT-PCR. The heat map shows mRNA levels relative to β-actin. Samples of human nasal (N=2) and lung tissue (N=8) were analyzed, besides the three cell lines (Calu-3, Vero E6 and HEK293T: N=2). We previously reported an expression analysis of all 18 human TTSPs in Calu-3 cells and human respiratory tissue (13). (B) Western blot analysis showing S expression and S1/S2 cleavage in HEK293T producer cells transfected with wild type (WT) or mutant forms of SARS-2-S, SARS-1-S or MERS-S.

### Cleavage of the SARS-2-S S1/S2 site is determined by the multibasic motif as well as the length of the cleavage loop

First, we examined processing of S0 into S1/S2, in HEK293T cells transfected with the WT and mutant S protein forms (Fig. 3B). The presence of a C-terminal V5-tag in the S proteins enabled western blot analysis with an anti-V5 antibody. The D614- and G614-forms of SARS-2-S showed a strong S2 band, indicating equally efficient S1/S2 cleavage by one or more proteases expressed in these cells (23–26). All three SARS-2-S^D614^ mutants bearing deletions in the S1/S2 loop showed virtually abrogated cleavage. The lack of cleavage for the ΔQTQTN mutant (which still possesses the multibasic furin motif but lacks preceding amino acids) indicates that not only the furin motif itself is critical for cleavage, but also the length of the loop presenting this motif. As expected (29), also WT MERS-S was efficiently cleaved, while its monobasic (monoR) cleavage site mutant was not processed. In contrast, SARS-1-S WT was barely cleaved, while proteolytic processing was efficient for the mutant containing the extended (Ext) S1/S2 cleavage loop of SARS-2-S, as anticipated (24).

### Loop deletion mutants of SARS-2-S show enhanced cathepsin-dependent entry, explaining their emergence in Vero cells

To evaluate how S1/S2 pre-cleavage impacts viral entry into Calu-3 or Vero E6 cells, we used WT and mutant pseudoviruses, produced at the optimal temperature established in the first part of this study. To fully discriminate the two S protein activation pathways, we included the protease inhibitors camostat and E64d (Fig. 4A). All three SARS-2-S^D614^ mutants bearing deletions in the S1/S2 cleavage loop showed markedly reduced (6-to 30-fold; P ≤ 0.009 versus WT) entry into Calu-3 cells (Fig. 4B, top left panel), in keeping with expectations (24). Entry was fully rescued when exogenous trypsin was added during Calu-3 cell transduction (Suppl. Fig. S2), indicating that the poor entry was due to a lack of S2’ cleavage by TTSPs, and not to inefficient receptor binding. Conversely, these three SARS-2-S loop deletion mutations resulted in 13-to 20-fold higher entry into Vero E6 cells, which depends on cathepsin L (Fig. 4B, top right panel). This explains why SARS-CoV-2 passaging in Vero E6 cells regularly leads to emergence of viruses bearing substitutions or deletions in the S1/S2 loop (30–36). For the SARS-1-S and MERS-S mutants, the data concurred with other reports. Namely, Calu-3 cell entry of SARS-1-S pseudovirus was unchanged when its S1/S2 sequence was exchanged for the extended loop of SARS-2-S, including the multibasic motif (24). Akin to SARS-2-S, mutant MERS-S pseudovirus bearing a monobasic (= non furin-cleavable) S1/S2 site showed dramatically reduced (71-fold) Calu-3 cell entry (28). On the other hand, entry into Vero E6 cells was more efficient (4-fold, P < 0.0001) for SARS-1-S-pseudovirus with WT protein (= short monobasic S1/S2 loop) than the mutant with extended multibasic loop. However, the opposite was seen for MERS-S, namely higher entry (3-fold; P = 0.0006) for WT (= furin-cleavable) than mutant (= monobasic) pseudovirus.

**Figure 4.**
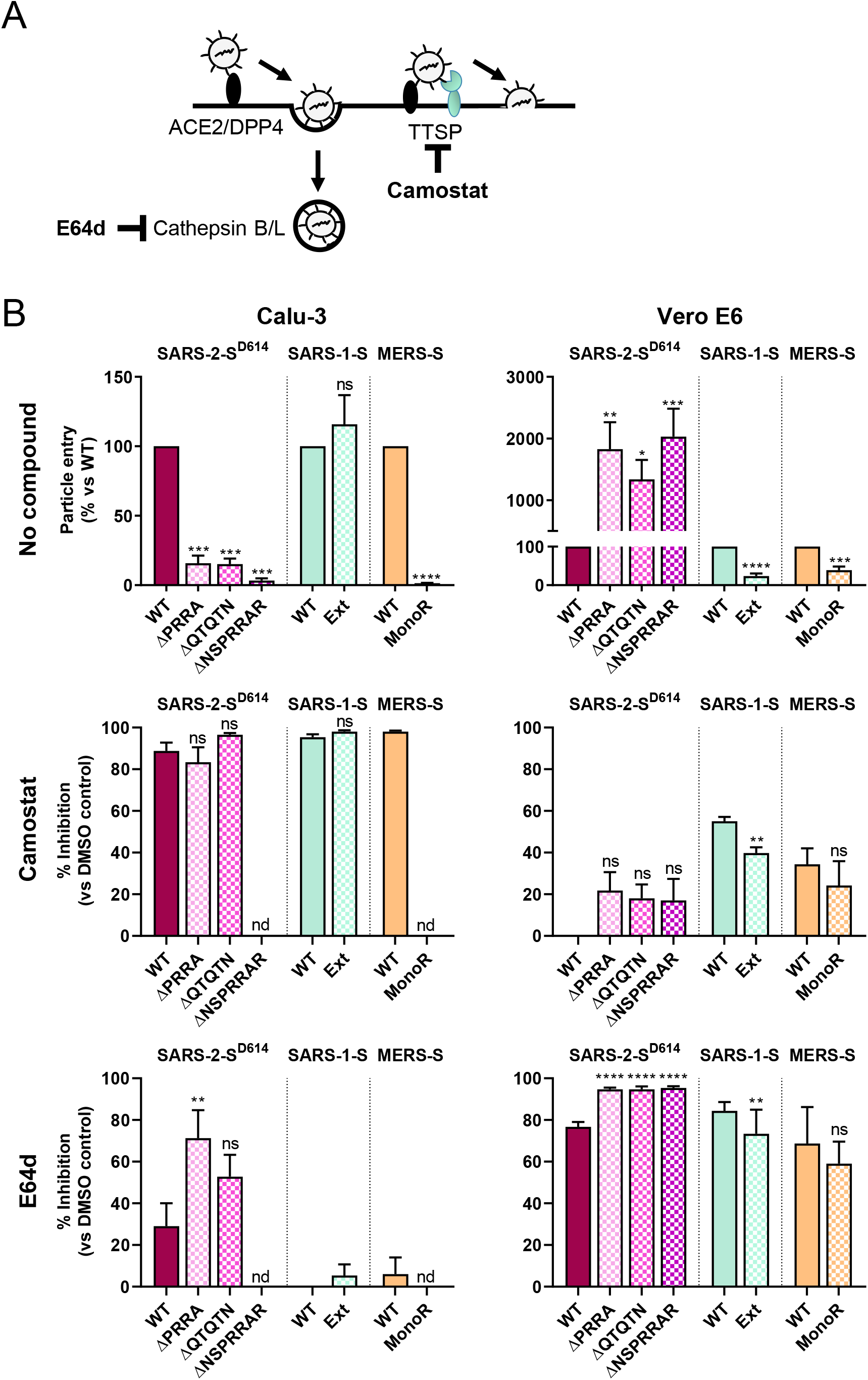
Entry of WT and S1/S2 mutant pseudoviruses in cells with different spike-activating proteases. (A) The broad serine protease inhibitor camostat prevents fusion activation by TTSPs like TMPRSS2, whereas E64d inhibits cathepsin B/L-mediated fusion after virus uptake by endocytosis. (B) The graphs show entry efficiency *(top panels)* or % inhibition relative to the DMSO solvent control, by 50 μM camostat *(middle panels)* or 50 μM E64d *(bottom panels);* in human airway epithelial Calu-3 cells *(left panels)*, which express TMPRSS2 and other TTSPs; and *(right panels)* cathepsin-rich Vero E6 cells. An ordinary one-way ANOVA with Dunnett’s correction was used to compare SARS-2 mutants versus WT and an unpaired two-tailed t-test was used to compare the WT and mutant forms of SARS-1 and MERS. ns, not significant, *, P ≤ 0.05; **, P ≤ 0.01; ***, P ≤ 0.001, ****, P ≤ 0.0001. Results are the mean ± SEM; N=3 with three independently produced stocks.

Camostat produced >80% inhibition of Calu-3 cell entry (Fig. 4B, middle left panel), corroborating that virus entry into these cells relies on serine proteases like TMPRSS2 (50). E64d (Fig. 4B, bottom left panel) had a small effect on the WT SARS-2-S particles but a 2-fold higher effect on the loop mutants, indicating that even in Calu-3 cells, the cathepsin route gets a boost when the pseudoparticles are not cleaved at S1/S2. In contrast, for all pseudoviruses, Vero E6 cell entry was highly sensitive (59-95% inhibition) to E64d but not camostat (Fig. 4B), confirming that S protein-driven entry into these cells is highly cathepsin L-dependent (16, 50, 51).

In summary, these results underline the previously proposed concept (23-26, 28) that SARS-2-S and MERS-S, but not SARS-1-S, require S1/S2 pre-cleavage in producer cells for subsequent TMPRSS2-dependent entry into Calu-3 cells. More importantly, we demonstrate that, for SARS-2-S, not only the integrity of the furin motif at the S1/S2 site but also the length of the loop harboring this cleavage site, are required to enable S1/S2 pre-cleavage. Mutants that cannot undergo this processing are boosted towards cathepsin B/L-mediated entry, explaining why substitutions or deletions in the S1/S2 cleavage loop commonly emerge during SARS-CoV-2 propagation in Vero E6 cells.

### Among all 18 TTSPs, TMPRSS2 and TMPRSS13 are the best activators of SARS-2-S

We next asked whether, besides TMPRSS2, other TTSPs can activate SARS-2-S for virus entry. The pseudoviruses were applied to TTSP-plus receptor-transfected HEK293T cells in the presence of E64d, to shut off the parallel cathepsin route (Fig. 5A). We first investigated activation of SARS-2-S by the 18 known human TTSPs or by furin (Fig. 5B). The most efficient activator was TMPRSS2, followed by TMPRSS13 that was only 3-fold less effective. Human airway trypsin-like protease (HAT) and furin were, respectively, 19- and 14-fold less active than TMPRSS2. Mutating the S1/S2 cleavage loop abrogated activation by TMPRSS2, TMPRSS13, HAT and furin (Fig. 5C), in accordance with the inability of these S proteins to mediate robust virus entry into Calu-3 cells. Next, we addressed whether these four proteases activate SARS-1-S, MERS-S and 229E-S (Fig. 5D). TMPRSS2 activated the S proteins of all four CoVs, in keeping with published data (50, 52-58). Intriguingly, TMPRSS13 enhanced entry driven by the S proteins of the highly virulent SARS-CoV-1, SARS-CoV-2 and MERS-CoV, but not the common cold virus HCoV-229E. MERS-S and 229E-S were both activated by HAT, with roughly the same efficiency as TMPRSS2. HAT proved effective on 229E-S and MERS-S, as reported earlier (56, 59). Finally, furin expression in the target cells gave weak activation of the four S proteins, which aligns with the report that extracellular furin can act at the stage of MERS-CoV entry (29). S1/S2 pre-cleavage was required for efficient TTSP activation of SARS-2-S and MERS-S, as evident from the much lower activation of the S1/S2 mutants compared to the WT (Fig. 5C and 5D). This effect may only apply to S proteins that naturally have a pre-cleavable S1/S2 site, since SARS-1-S showed equal activation by TTSPs whether it was or was not pre-cleaved (compare WT and Ext mutant in Fig. 5D).

**Figure 5.**
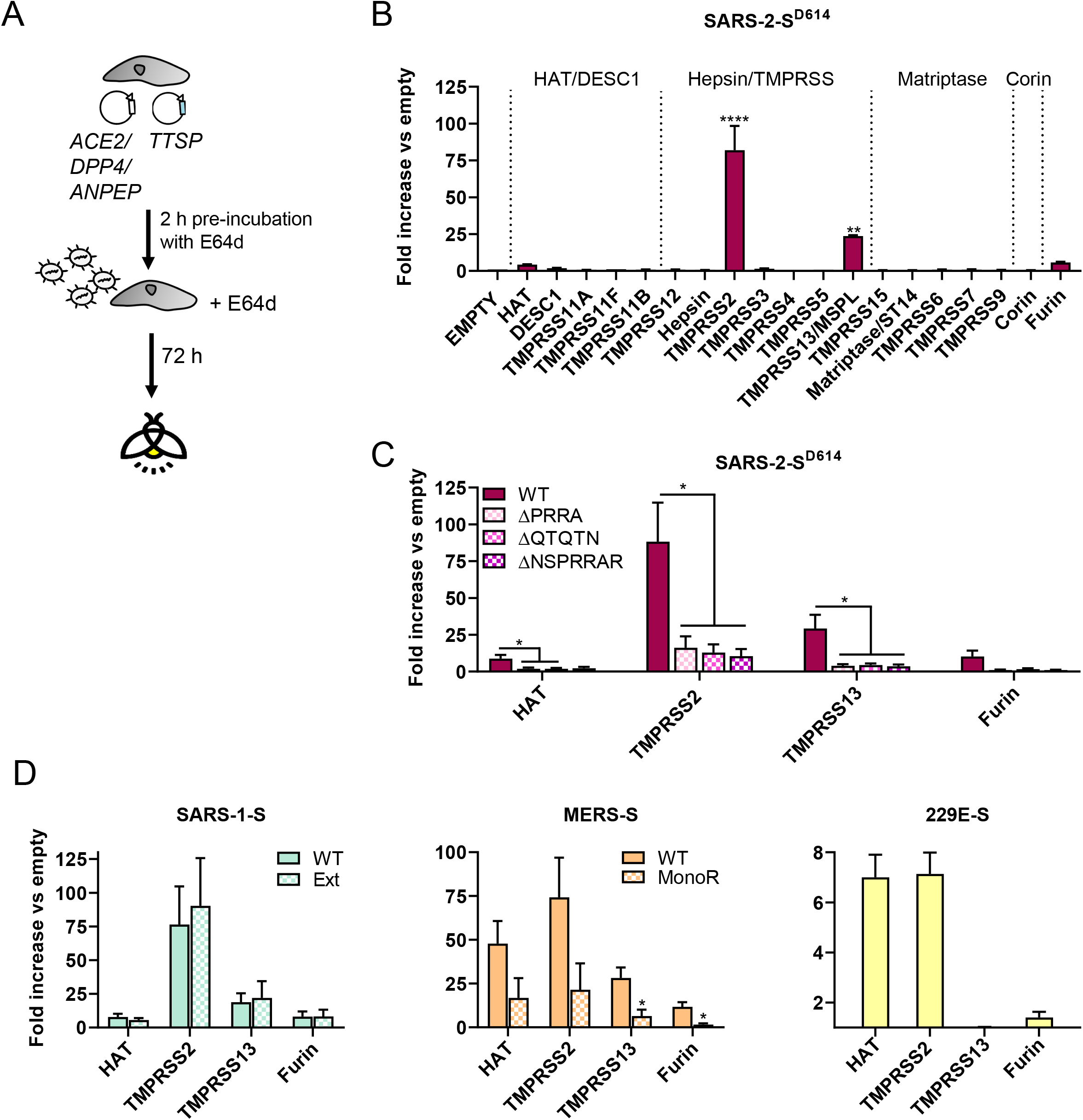
Activation of virus entry by different human TTSPs. (A) Experiment set-up. One day before transduction, HEK293T target cells were transfected with the appropriate receptor and one of the TTSPs. To block the cathepsin route, E64d was added at 2 h before and during transduction. (B) SARS-2-S activating capacity of the 18 human TTSPs. At the top of the graph, the four TTSP subfamilies are indicated. (C, D) The four TTSPs that proved active in panel B were evaluated for activation of wild-type and mutant forms of SARS-2-S (panel C), or SARS-1-S, MERS-S and 229E-S (panel D). An ordinary one-way ANOVA with Dunnett’s correction was used to compare SARS-2 mutants versus WT and an unpaired two-tailed t-test was used to compare the WT and mutant forms of SARS-1 and MERS. *, P ≤ 0.05; **, P ≤ 0.01; ****, P ≤ 0.0001. Results are the mean ± SEM; N=3 with three independently produced stocks).

To summarize, these results corroborate TMPRSS2 as an efficient and broad S protein activator, and identify TMPRSS13 as a novel activator of highly pathogenic CoVs.

### The D614G change increases SARS-CoV-2 spike stability, reduces its reliance on 33°C and boosts entry via the cathepsin route

Finally, we investigated the impact of the D614G mutation on the temperature and protease dependency of the SARS-2-S protein. We first assessed pseudovirion thermostability, since one study indicated that the D614G mutation generates a more stable trimeric spike protein by removing a destabilizing inter-protomer interaction (44, 45). Also, since the S^D614^ protein may more easily shed the S1 subunit (45), we included our SARS-2-S mutants bearing unprimed spikes due to S1/S2 loop deletions. Finally, we included the SARS-1-S, MERS-S and 229E-S pseudovirions. In order to assess S protein stability, these different pseudotypes were incubated for 1 h at varying temperatures (range: 33 to 41°C, and 4°C for the control) and then tested for infectivity in HEK293T cells expressing receptor and TMPRSS2 (Fig. 6A). SARS-2-S^D614^ had comparable stability as SARS-1-S and 229E-S, while MERS-S appeared slightly more stable (Fig. 6B). On the other hand, the thermostability of SARS-2-S^D614^ was markedly increased when it was not pre-cleaved at S1/S2 (see the three loop deletion mutants in Fig. 6C). Importantly, the same stabilizing effect was achieved by introducing the D614G substitution (Fig. 6C).

**Figure 6.**
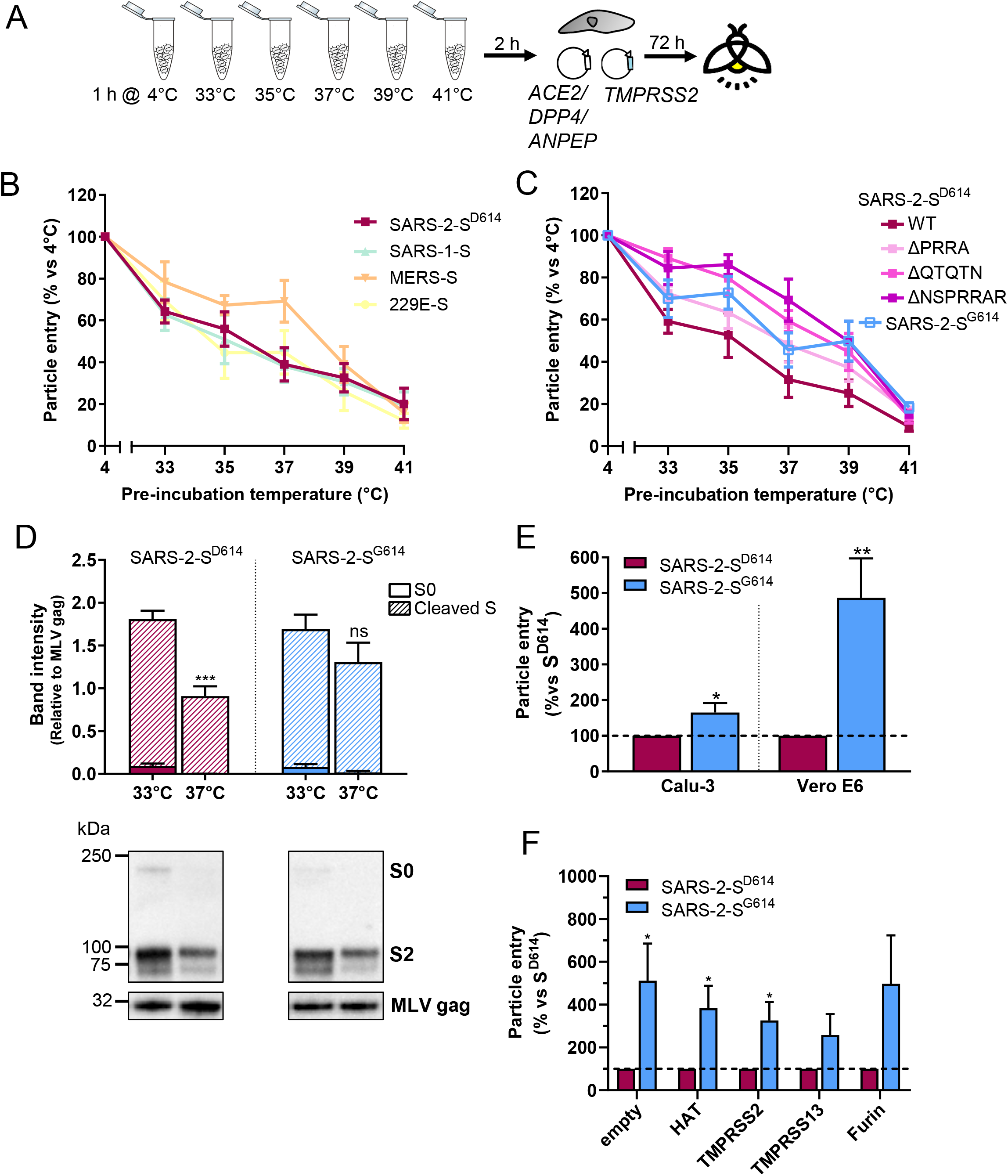
The SARS-2-S D614G change increases stability and entry via the cathepsin route, and reduces the preference for 33°C. (A) Thermostability analysis. The pseudoparticles were incubated at the indicated temperatures for 1 h, followed by 2 h entry into HEK293T target cells and luminescence reading after 72 h. Results are expressed as transduction efficiency, relative to the condition incubated at 4°C (mean ± SEM, N=3). (B) analysis of the four CoV pseudotypes; (C) comparison of SARS-2-S^D614^, the three S1/S2 loop mutants, and the SARS-2-S^G614^ mutant. (D) Pseudovirions carrying SARS-2-S^D614^ or S^G614^ were produced at 33°C or 37°C, then pelleted to determine their S content by western blot. The graphs show the S0 and S2 band intensities, normalized to the band of MLV-gag (mean ± SEM, N=3). (E) Cell entry efficiency (mean ± SEM, N=3) of the two pseudovirus variants in Calu-3 and Vero E6. (F) Particle entry in HEK293T cells transfected with an empty plasmid (*empty*) or one of the TTSPs. In all panels: ns, not significant; *, P ≤ 0.05; **, P ≤ 0.01; ***, P ≤ 0.001, unpaired two-tailed t-test.

Secondly, the SARS-2-S^G614^ variant was less dependent on 33°C to achieve high protein levels in released pseudovirions (Fig. 6D). The level of the S^G614^ protein was not significantly higher when the particles were produced at 33°C instead of 37°C, while S^D614^ reached a significantly (P=0.0002) higher level at 33°C (Fig. 6D). The two variants showed no difference in terms of S1/S2 cleavage efficiency, and both generated pseudovirions with fully S1/S2 pre-cleaved spikes. This contradicts another study (46) but agrees with three other reports (40, 42, 44). Last but not least, compared to S^D614^, S^G614^ pseudovirus showed superior entry into Calu-3 and Vero E6 cells (Fig. 6E). The enhancement was particularly prominent in Vero E6 cells, suggesting that the D614G mutation mainly promotes cathepsin B/L-dependent entry. In line with this, the S^G614^ pseudovirus proved independent of TTSP activation to enter into HEK293T cells. Compared to S^D614^ pseudovirus, it entered 3-to 5-fold more efficiently, but this gain was similar whether the cells were transfected with a TTSP-expression or empty plasmid (Fig. 6F).

To conclude, our results indicate that the D614G mutation generates a more stable SARS-2-S protein, which might offer the S^G614^ variant a growth advantage. The mutation causes a shift towards cathepsin-mediated entry and reduces the spike’s reliance on 33°C to achieve high S protein levels in pseudoparticles.

## DISCUSSION

In this study, we recognized two features of the SARS-CoV-2 spike protein, i.e. compatibility with the varying temperature in the human respiratory tract and a well-tuned protease activation mechanism, which may well be two determinants for the high transmissibility and virulence of this virus. Our approach to include pseudoviruses with the spikes of SARS-CoV-1, MERS-CoV and HCoV-229E, offered the possibility to notice analogies and interpret our findings from a broader perspective.

First of all, we unveiled a distinct temperature preference for these different CoV spike proteins, that precisely matches the predilection of each virus for the upper or lower respiratory tract. SARS-CoV-2, but not SARS-CoV-1, replicates abundantly in the nose (~30-32°C) and upper airways (1, 2), a driving factor behind its efficient transmission. Both viruses, as well as MERS-CoV, also replicate in the lungs (37°C) to produce severe pathology. Intriguingly, we found that SARS-1-S and, to a lesser extent, MERS-S favour 37°C, while SARS-2-S is compatible with 37°C, yet prefers 33°C. This aligns with viral behaviour in cell culture, since SARS-CoV-2, but not SARS-CoV-1, was shown to prefer 33°C to 37°C (48). We observed the same for the common cold HCoV-229E, which prefers a cooler temperature for virus replication and for which 37°C proved detrimental when generating pseudoparticles bearing its spike. This airway temperature preference is a property of the coronavirus spike glycoprotein, but also the influenza virus hemagglutinin (13), suggesting that it might be a commonality for human respiratory viruses. As to the biochemical basis, a cooler temperature likely ensures lower conformational flexibility during S glycoprotein synthesis or transport (60), and avoidance of unstable spike conformers. For SARS-2-S, the 33°C preference proved more pronounced for the S^D614^ than the S^G614^ variant. The S^D614^ form may require 33°C for productive folding and transport, while the S^G614^ protein appears less demanding thanks to its higher stability, as evident from our thermostability analysis. Our findings generated with pseudotyped virus are supported by data with authentic viruses, since the SARS-CoV-2-S^D614^ and -S^G614^ variants exhibited comparable replication in human nasal cells at 33°C, but the S^G614^ variant reached higher titers in bronchial epithelial cells at 37°C and 39°C (41).

Secondly, we revealed which of the 18 human TTSPs, of which many are present in human respiratory tissue (13), can activate SARS-2-S for virus entry. We confirmed that TMPRSS2 is an efficient and broad CoV spike activator. HAT proved effective on 229E-S and MERS-S, however it was less active on SARS-1-S and SARS-2-S, as reported earlier (54, 56, 59, 61).The most intriguing result regards TMPRSS13, which we identified as a second potent activator of SARS-1-S, SARS-2-S and MERS-S, but not 229E-S. Since this protease prefers cleavage sites with a second basic residue at positions P2 or P4 (62), it plausibly recognizes the S2’ site at the KR motif of SARS-1-S and SARS-2-S, and the RSAR motif of MERS-S. Such a motif is missing in the predicted S2’ sites of all common cold CoVs (21). This raises the hypothesis that TMPRSS13 cleavability might be a CoV virulence factor, in line with the observation that this protease activates the hemagglutinin of some highly pathogenic avian influenza A viruses (62). TMPRSS13 is highly expressed in different cell types of the human respiratory tract (61, 63) and is also present in Calu-3 cells (13). It is important to note that SARS-2-S cleavability by other TTSPs than TMPRSS2 does not compromise clinical evaluation of camostat against COVID-19 (ClinicalTrials.gov identifiers: NCT04455815, NCT04321096, NCT04353284, NCT04355052, NCT04374019), since this molecule is a broad TTSP inhibitor (61).

Thirdly, while SARS-1-S did not need S1/S2 pre-cleavage, this processing proved crucial for TTSP-activation of SARS-2-S, presumably by provoking a conformational change that facilitates subsequent S2’ cleavage (24, 28, 64). Our data with SARS-2-S loop mutants show that the extended loop length and furin motif are equally important to achieve S1/S2 processing. Besides furin (or related proprotein convertases), also cathepsin B/L in the secretory pathway (65) might possibly perform this pre-cleavage during S protein trafficking or during viral egress via a recently discovered lysosome-exocytic pathway (66). All three enzymes were shown to cleave the S1/S2 sequence in an enzymatic assay (27). When unprimed, SARS-2-S pseudoviruses are strongly boosted towards the cathepsin B/L route. This likely explains the replication advantage of loop-deletion SARS-CoV-2 mutants in cathepsin L-rich Vero E6 cells (30–36). Whereas the S1/S2 primed state predisposes to S2’ activation by TTSPs, the non-covalently linked S1/S2 form is less stable and a plausible disadvantage for endosomal entry. During virus traffic from the cell membrane to late endosomes [which takes up to 1 h (67)], S1 and S2 must remain associated under gradually more acidic pH conditions. The S1/S2 loop mutant (= unprimed) virions circumvent this problem, since they are only cleaved by cathepsins after reaching acidic endosomes. The superior stability of loop mutants over WT SARS-2-S^D614^ was also evident in our thermostability experiments.

Strikingly, also the more stable S^G614^ variant proved to be boosted for Vero E6 cell entry, despite being pre-cleaved. Its higher stability concurs with lower S1 shedding (45), a clear advantage for endosomal entry. Improved stability may be the reason why SARS-CoV-2-S^D614^ was superseded by the S^G614^ variant within a few months of circulation in humans (43). Animal studies showed that the SARS-CoV-2-S^G614^ variant has higher fitness and transmissibility (40-42). This is reminiscent of influenza virus, for which the link between hemagglutinin stability and transmissibility is well established (13, 68). The relatively low stability of S^D614^ could be attributed to an unfavorable interprotomer contact that is not present in S^G614^ (44, 45). Besides, the superior cell entry of S^G614^ pseudovirus, seen in this and several other studies (42–47), was rationalized by structural evidence that the D614G substitution leads to a more open receptor binding conformation (44). This can explain increased entry of the S^G614^ variant via the TTSP pathway in Calu-3 cells (40, 44). However, we found that this gain was quite modest, while entry via the endosomal pathway proved much more enhanced. This indicates that the S^G614^ virus is more effective at using the two entry routes, which may broaden its cell tropism. In cells lacking appropriate surface proteases, endosomal entry might provide an alternative pathway. Cathepsin B/L-mediated CoV entry may thus have higher in vivo relevance than often assumed. Its role might be underestimated when Calu-3 cells are used, since they express low levels of cathepsin B and cathepsin L. Both cathepsins are expressed at high levels in several types of airway epithelial cells (69), in line with our analysis on nasal and lung tissue samples.

Of note, endosomal entry exposes the virus to endosomal interferon-induced transmembrane proteins (IFITMs), which may either act as virus restriction factors (70) or, in sharp contrast, be hijacked by SARS-CoV-2, as recently demonstrated (71). Here, temperature might again be a key player, as innate immune activation proved to be greatly diminished at 33°C compared to 37°C (48, 72). In our experiments, unprimed (= S1/S2 mutant) MERS-S pseudovirus differed from the SARS-2-S counterparts, in not showing higher entry into Vero E6 cells, which is in line with another study (28). This might suggest that the MERS spike is particularly sensitive to IFITM control or highly unstable at endosomal pH. Yet, these hypotheses definitely need validation with live virus in appropriate human airway models.

To conclude, we revealed mechanisms whereby the coronavirus spike protein is adjusted to match the temperature and protease conditions of the human airways. This insight will help to better comprehend coronavirus-host interaction and adaptation, and, in short term, will be highly valuable to understand the behaviour of emerging spike mutants of SARS-CoV-2.

## MATERIALS AND METHODS

### Ethics statement

Lung tissue samples from eight different healthy donors and nasal tissue samples from one healthy donor and one patient with chronic rhinosinusitis with nasal polyps were obtained under the approval of the ethical committee from the University Hospital Leuven (UZ Leuven Biobanking S51577 and S59864). All patients were adult and provided written informed consent.

### Cells, media and compounds

Calu-3 (ATCC HTB-55) cells were grown in minimum essential medium (MEM) supplemented with 10% fetal calf serum (FCS), 0.1 mM non-essential amino acids, 2 mM L-glutamine, and 10 mM HEPES. HEK293T cells (HCL4517; Thermo Fisher Scientific) and Vero E6 (ATCC CRL-1586) were grown in Dulbecco’s modified Eagle’s medium (DMEM) supplemented with 10% FCS, 1 mM sodium pyruvate, 0.075% sodium bicarbonate and (only for Vero E6) 0.1 mM non-essential amino acids. Unless stated otherwise, all cell incubations were done at 37°C.

The transduction medium consisted of DMEM supplemented with 2% FCS, 0.1 mM non-essential amino acids, 1 mM sodium pyruvate, 0.075% sodium bicarbonate, 100 U/mL of penicillin and 0.1 mg/mL of streptomycin. Camostat mesylate was purchased from Sigma-Aldrich, whereas E64d and chloromethylketone (dec-RVKR-CMK) were from Enzo Life Sciences.

### Analysis of receptor and protease gene expression in cells, and human lung and nasal tissue

Total RNA was extracted using a ReliaPrep RNA Cell Miniprep System (Promega), and 0.5 μg of RNA was converted to cDNA with a high-capacity cDNA reverse transcription kit (Thermo Fisher Scientific). BRYT Green dye-based quantitative PCR (qPCR) was performed with GoTaq qPCR Master Mix (Promega) and intron-spanning primer pairs (see Suppl. Table S4 for a list of all primers) in an ABI 7500 Fast real-time PCR system (Applied Biosciences). Expression data were normalized to housekeeping gene *ACTB*.

### Plasmids

The expression plasmids for HAT and DESC1 were reported before (73). The plasmids for the other 16 TTSPs and furin were purchased from GenScript and described earlier (13). To express the CoV receptors, we used plasmids encoding human ACE2 (74); human DPP4 (75) and human APN (56).

The plasmids to express SARS-1-S and MERS-S with a C-terminal V5-tag were already reported (76, 77). To create the SARS-2-S-V5 expression plasmid, we used a starting plasmid carrying a codon-optimized SARS-2-S coding sequence (early pandemic D614 variant; GenBank ID MN908947.3) that was generously provided by K. Dallmeier (Leuven, Belgium) (78). A C-terminal V5-tag was added and the construct was subcloned into the pCAGGS vector using the NEBuilder HiFi DNA Assembly kit (New England Biolabs). Likewise, a V5 tag was introduced into a pCAGGS-based plasmid encoding 229E-S (79). Mutations in the S coding sequence were introduced via PCR with overlapping primers, and inserted into pCAGGS using the NEBuilder HiFi DNA Assembly kit. All plasmids were subjected to sequencing analysis to verify the presence of the desired mutations and absence of any unwanted mutations.

### Production of MLV-S pseudoviruses and transduction experiments

The method to produce firefly luciferase (fLuc)-expressing MLV pseudovirus carrying CoV S-protein, was previously described (80). In brief, HEK293T cells seeded in 6-well plates, were transfected using Lipofectamine® 2000 (Life Technologies), with a mixture of plasmids encoding MLV gag-pol, the fLuc reporter and V5-tagged S-protein. At 4 h post transfection, the medium was replaced by medium with 2% FCS. Pseudoparticle production was done at 33°C or 37°C, as specified in the Figure legends. At 48 h, the pseudovirus-containing supernatants were harvested, clarified by centrifugation and stored at −80°C.

For transduction (always performed at 37°C), Calu-3 or Vero E6 cells were seeded in white 96-well plates and one day later exposed to 100 μl virus stock. In the case of HEK293T, the cells were first transfected with the receptor- and TTSP-expression plasmids, at 24 h before transduction. In some experiments, protease inhibitors, i.e. camostat mesylate, E64d, chloromethylketone (all at 50 μM) or 1% DMSO (solvent control), were added at 2 h before pseudovirus transduction. At 6 h after transduction, pseudovirus and compounds were removed and fresh medium was added. Three days later, fLuc activity was measured using a luciferase assay system kit and GloMax^®^ Navigator Microplate Luminometer (both from Promega).

To assess thermostability of the pseudoparticles, they were incubated for 1 h in tubes, at a temperature of 33, 35, 37, 39 or 41°C, or at 4°C included as control. They were then transduced into receptor- and TMPRSS2-transfected HEK293T cells. Two hours later, the transduction medium was replaced by complete growth medium.

### Western blot analysis of S protein expression and incorporation into pseudoparticles

To analyze S protein expression, the plasmids encoding a V5-tagged S protein were transfected into HEK293T cells, using Lipofectamine® 2000. Four hours later, the medium was replaced and the cells were incubated for another 48 h. Next, the cells were washed once with PBS and lysed in RIPA buffer supplemented with protease inhibitor cocktail (both from Thermo Fisher Scientific). The lysates were boiled for 5 min at 95°C in 1x XT sample buffer containing 1x XT reducing agent (both from Bio-Rad) and resolved on 4-12% Bis-Tris XT precast gels (Bio-Rad). The proteins were transferred to polyvinylidene difluoride membranes (Bio-Rad), blocked with 5% low-fat milk solution, and probed for 1 h with primary antibody followed by 45 min with secondary antibody. Bands were detected using SuperSignal West Femto Maximum Sensitivity Substrate (Thermo Fisher Scientific) and a ChemiDoc XRS+ system (Bio-Rad). The primary antibodies were mouse anti-V5 tag [Invitrogen, R960-25, 1:2000 (SARS-1-S, MERS-S and SARS-2-S) or 1:5000 (229E-S)] and mouse anti-β-actin (Sigma-Aldrich, A5447, 1:5000). As secondary antibody, we used a peroxidase-coupled goat anti-mouse antibody (Dako, P0447, 1:5000).

For analysis of S protein incorporation into pseudoparticles, a volume of 600 μl of S-pseudotyped MLV virus was loaded onto a 20% (w/v) sucrose cushion (volume 50 μl) and subjected to high-speed centrifugation (25,000 g for 120 min at 4°C). Thereafter, 630 μl of supernatant was removed and the residual volume was mixed with 30 μl loading dye mastermix, consisting of RIPA buffer supplemented with protease inhibitor cocktail (both from Thermo Fisher Scientific) and 1x XT sample buffer containing 1x XT reducing agent (both from Bio-Rad). The samples were heated for 5 min at 95°C and subjected to SDS-PAGE and immunoblotting, as above, with mouse anti-V5 tag (Invitrogen, R960-25) and mouse anti-MLV p30 antibody (Abcam, ab130757) as primary antibodies.

### Statistical analysis

Statistical analysis was performed using GraphPad Prism (version 8.4.3). An ordinary one-way ANOVA with Dunnett’s correction was performed when comparing multiple groups, while an unpaired two-tailed t-test was used when comparing only two groups. P ≤ 0.05 was considered significant. Statistical significance is reported as: *, P ≤ 0.05; **, P ≤ 0.01; ***, P ≤ 0.001; ****, P ≤ 0.0001.

## ACKNOWLEDGEM ENTS

This research work was supported by funding from the European Union’s Innovative Medicines Initiative (IMI) under Grant Agreement 101005077 [Corona Accelerated R&D in Europe (CARE) project], and from Fundació La Marató de TV3, Spain (Project No. 201832-30). The authors wish to thank Julie Vandeput, Joren Stroobants and Lisa Rectem for technical assistance; Kai Dallmeier and Lorena Sanchez Felipe for providing the SARS-2-S starting plasmid; Piet Maes for the Vero E6 cells and Bart Vanaudenaerde for providing human lung tissue samples.

## Author Contributions

M.L., A.S, and L.N. designed research; M.L., V.R., and R.V.B. performed research; M.L., A.S., and L.N. analyzed data; K.M., and S.P. contributed materials; and M.L., A.S., S.P., and L.N. wrote the paper.

## Classification

Biological sciences, Microbiology

## Supplementary Information

### This PDF file includes

Supplementary Tables S1 to S4

Supplementary Figures S1 to S2

**Supplementary Table S1.**
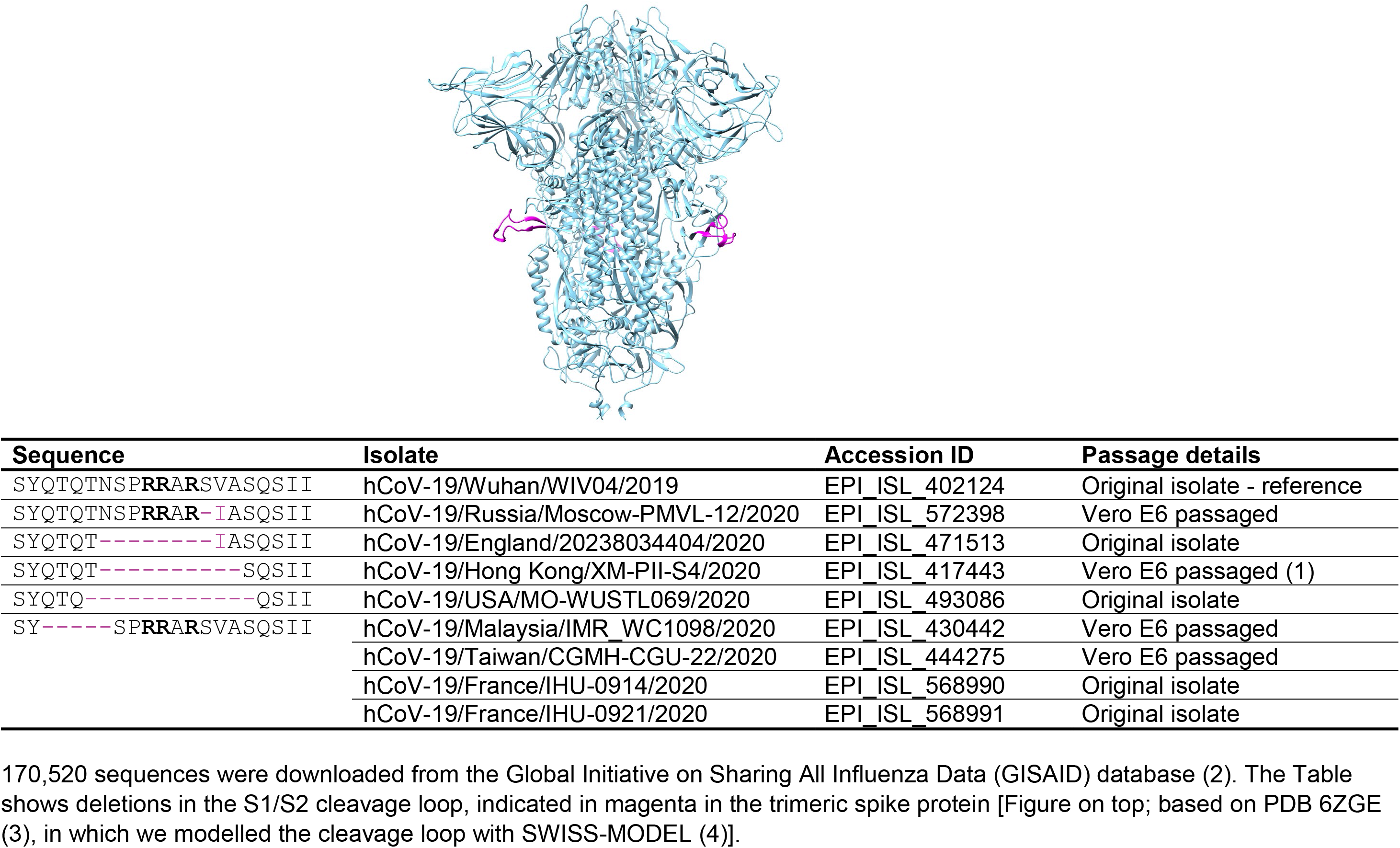
Deletions in the S1/S2 cleavage loop, observed in the GISAID database.

**Supplementary Table S2.**
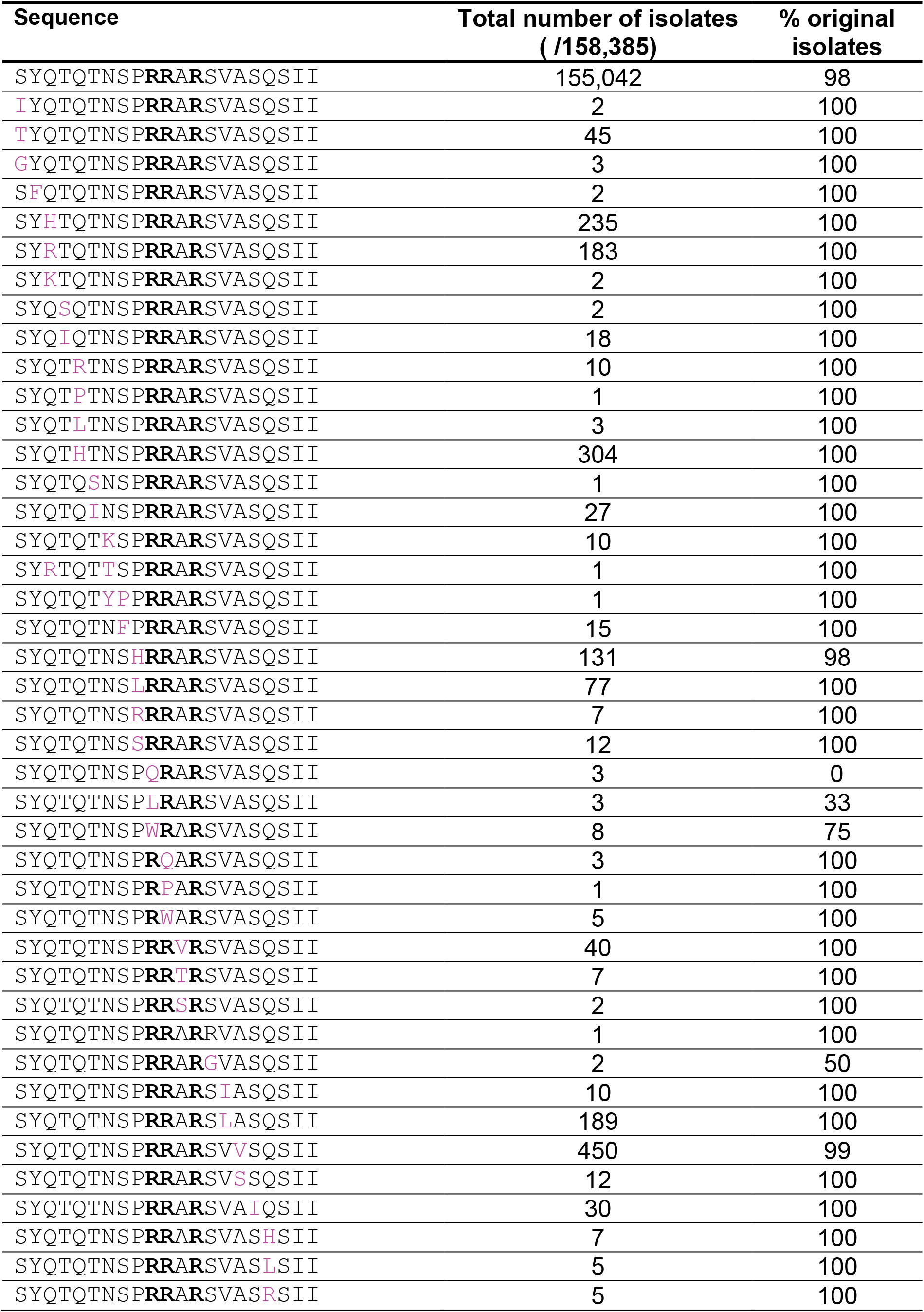
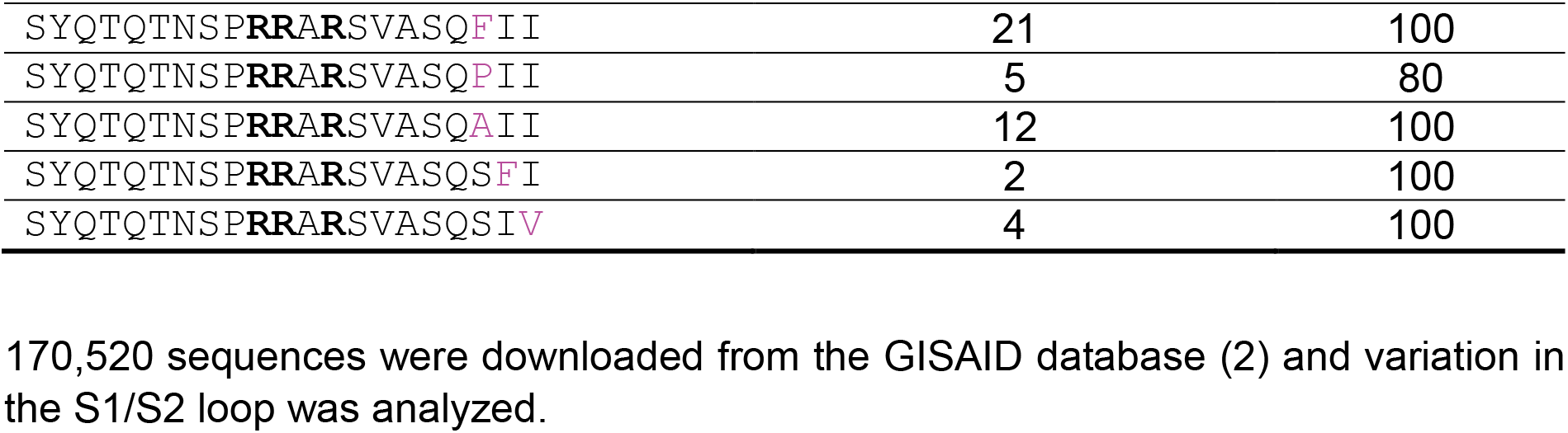
Substitutions in the S1/S2 cleavage loop, observed in the GISAID database.

**Supplementary Table S3.**
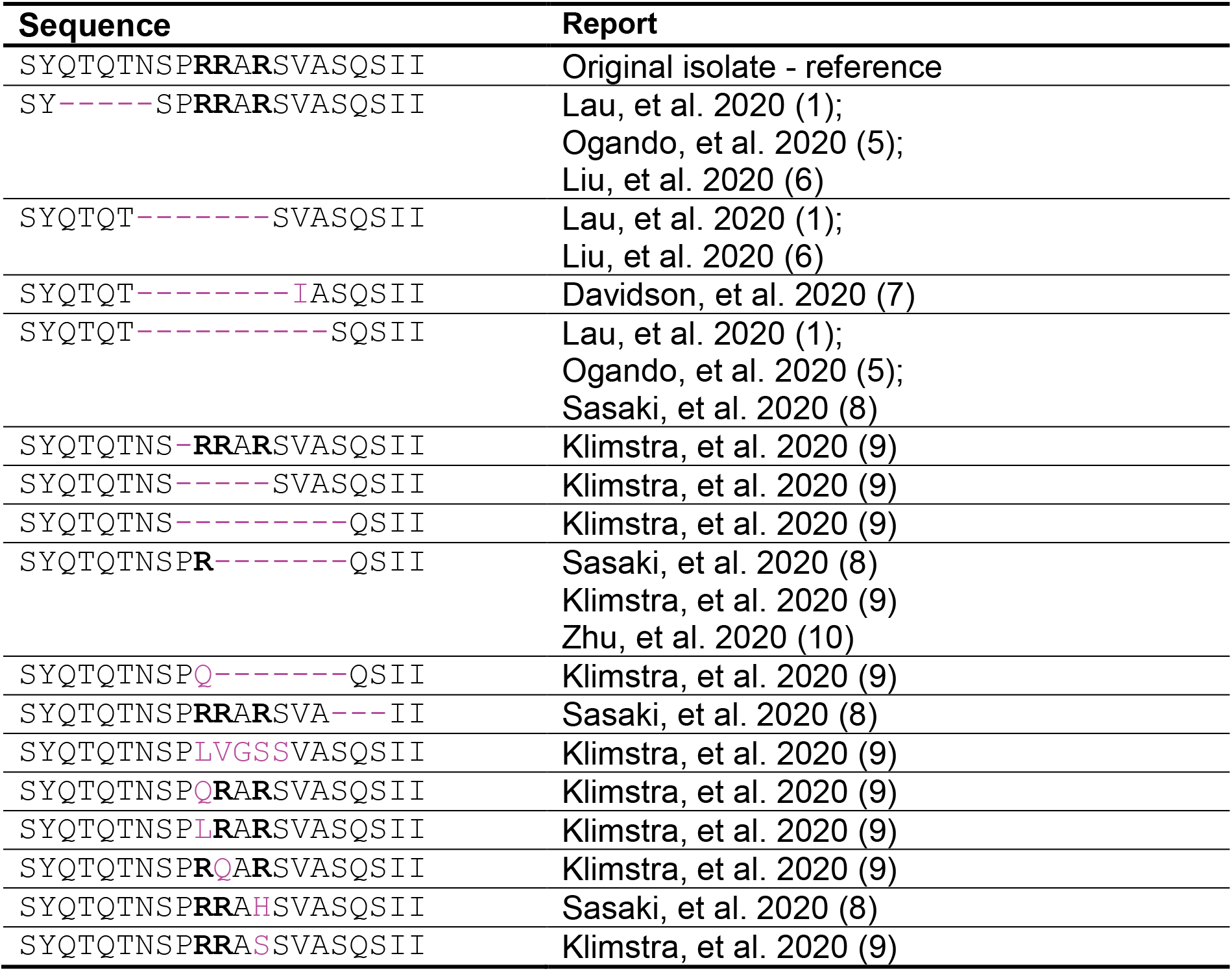
Deletions and substitutions in the S1/S2 cleavage loop, observed after passaging in cell culture.

**Supplementary Table S4.**
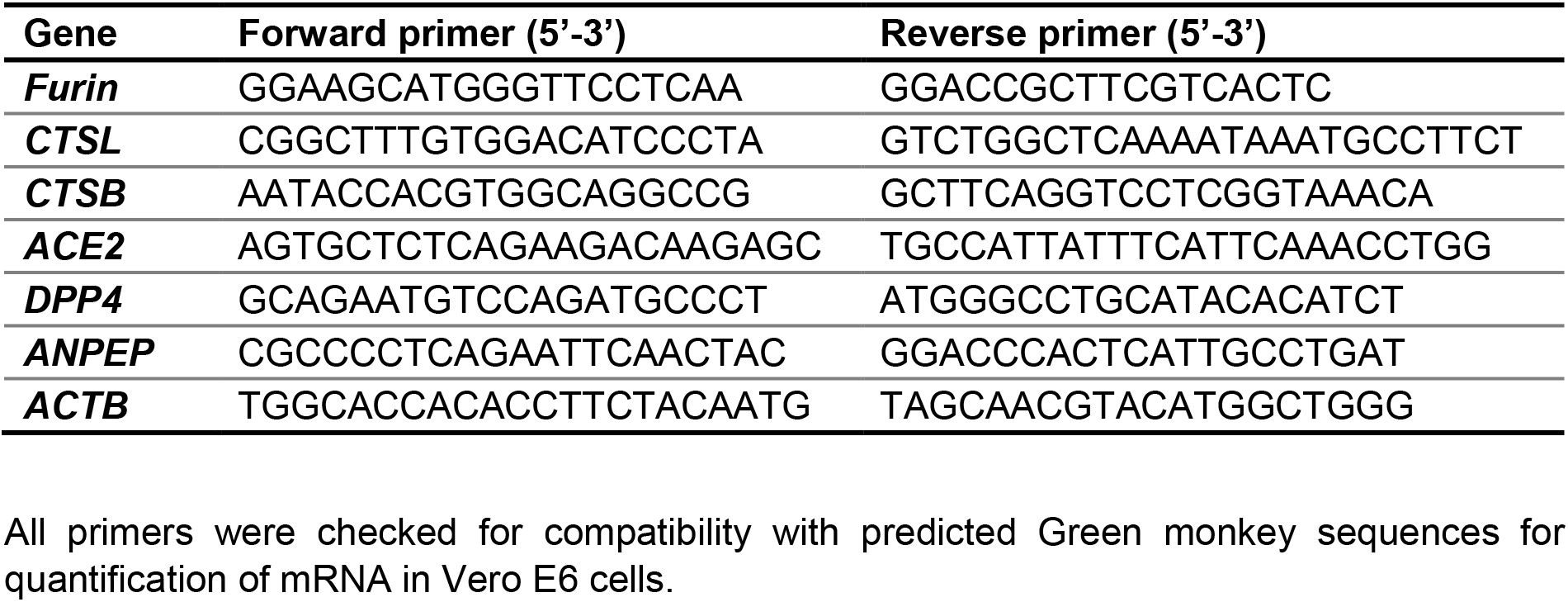
Primers for RT-qPCR quantification.

**Figure S1.**
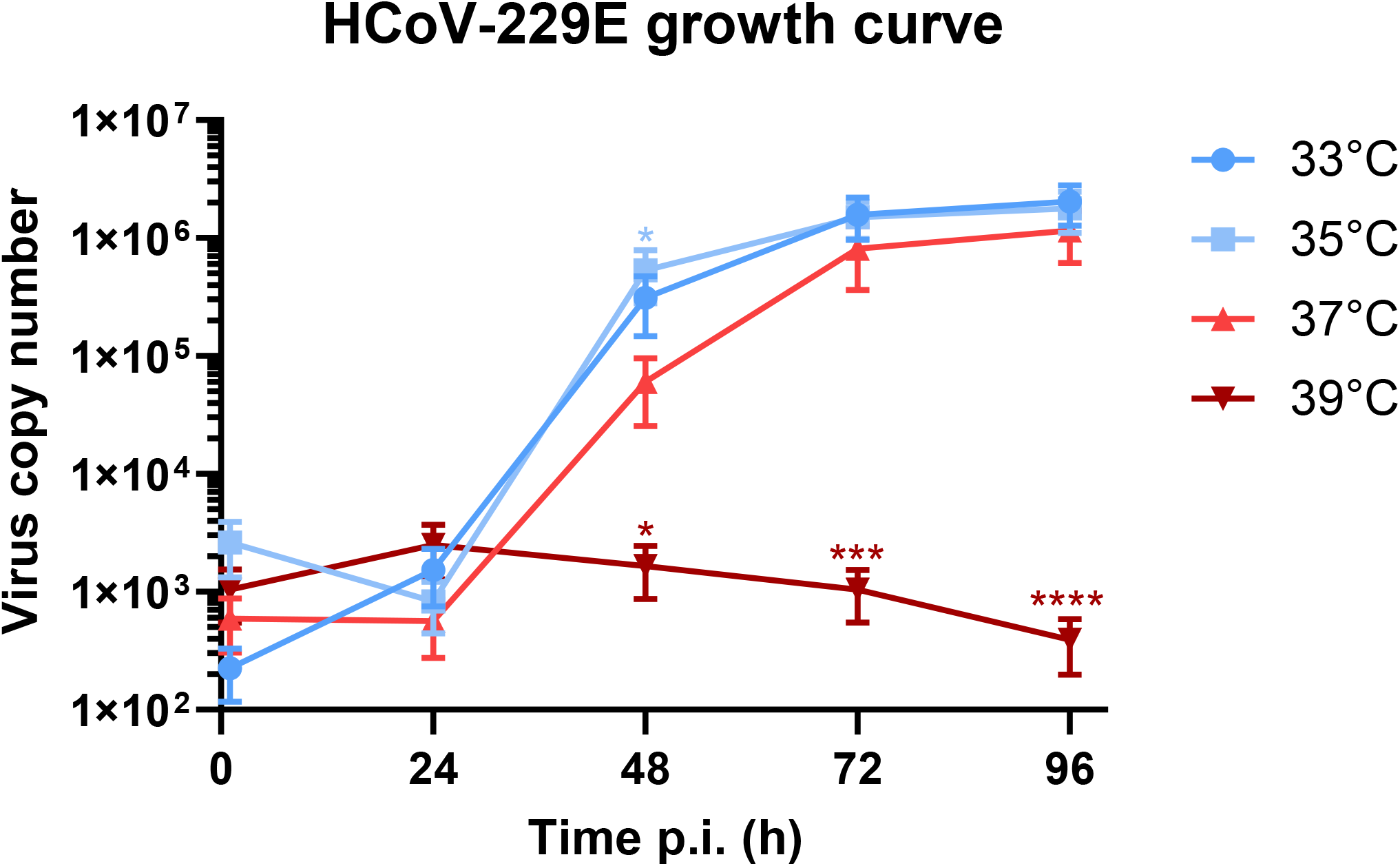
HCoV-229E shows temperature-dependent replication with a preference for 33 and 35°C. Human embryonic lung (HEL) fibroblast cells were infected with HCoV-229E at 100xCCID_50_ (determined at 37°C). The viral RNA copy number in the supernatant was determined at 1,24, 48, 72 and 96 h post infection (p.i.) by RT-qPCR with HCoV-229E N-gene specific primers and probe, as described elsewhere (11). An ordinary one-way ANOVA with Dunnett’s correction was used to compare the virus copy number at 33, 35 and 39°C vs 37°C. *, P ≤ 0.05; ***, P ≤ 0.001, ****, P ≤ 0.0001, ordinary one-way ANOVA with Dunnett’s correction.

**Figure S2.**
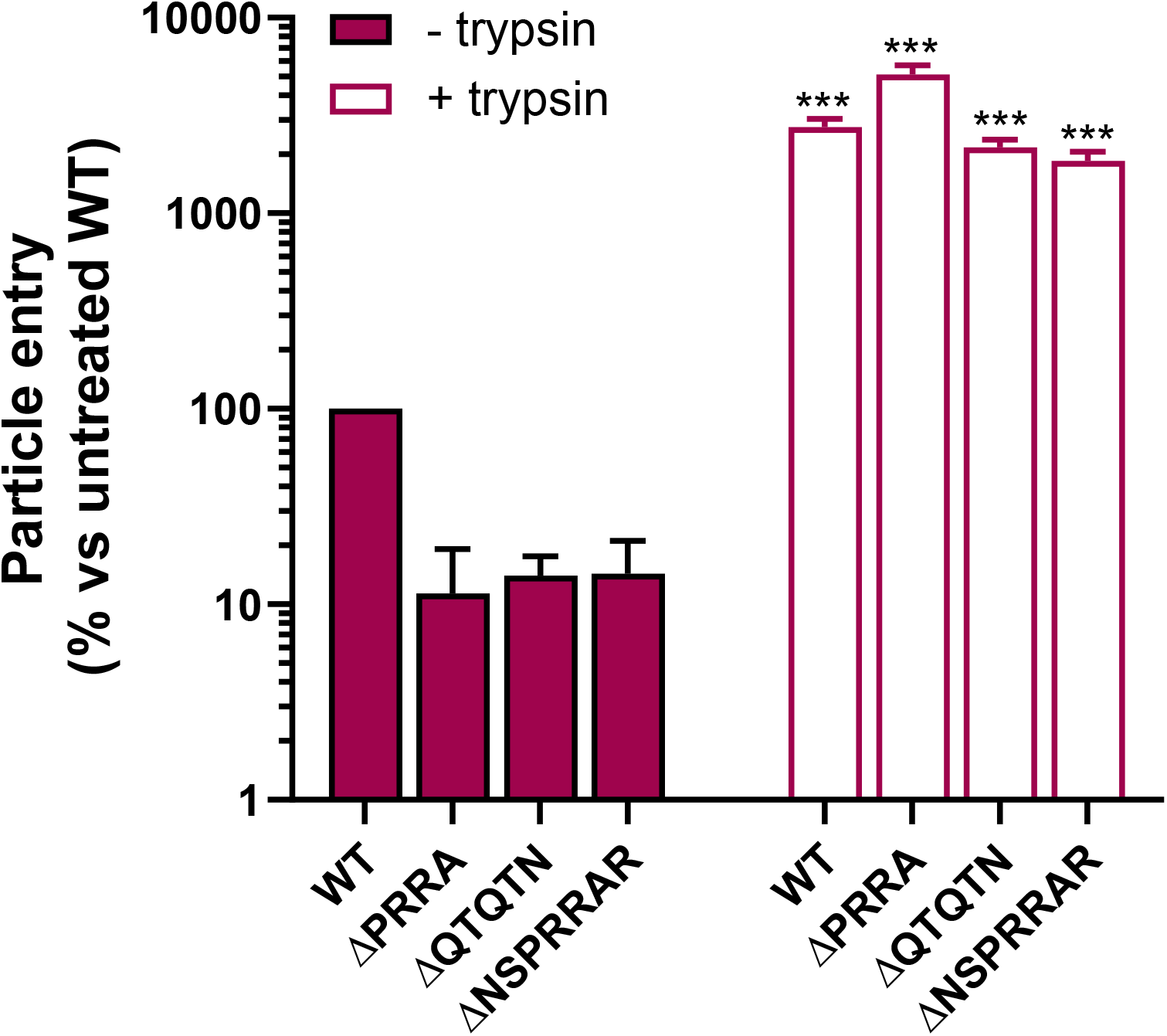
The Calu-3 cell entry defect of S1/S2 loop mutant SARS-2 pseudoviruses is restored when exogenous trypsin is added. Particles were allowed to bind for 1 h at 4°C after which unbound particles were removed and DMEM as such with 10 μg/ml TPCK-trypsin was added. After 2 h at 37°C, the medium was changed again to complete medium with 10% FCS. Results are the mean ± SEM; N=3. Each trypsin-treated condition was compared to untreated by an unpaired two-tailed t-test. ***, P ≤ 0.001.

